# SARS-COV-2 Spike Protein Fragment eases Amyloidogenesis of α-Synuclein

**DOI:** 10.1101/2023.05.06.539715

**Authors:** Andrew D. Chesney, Buddhadev Maiti, Ulrich H. E. Hansmann

## Abstract

Parkinson’s Disease is accompanied by presence of amyloids in the brain formed of α-synuclein chains. Correlation between COVID-19 and the onset of Parkinson’s disease let to the idea that amyloidogenic segments in SARS-COV-2 proteins can induce aggregation of α-synuclein. Using molecular dynamic simulations, we show that the fragment FKNIDGYFKI of the spike protein, which is unique for SARS-COV-2, shifts preferentially the ensemble of α-synuclein monomer towards rod-like fibril seeding conformations, and at the same time stabilizes differentially this polymorph over the competing twister-like structure. Our results are compared with earlier work relying on a different protein fragment that is not specific for SARS-COV-2.

## I. INTRODUCTION

Several neurodegenerative diseases are correlated with presence of protein aggregates in the brain, amyloids, that are characterized by a cross-beta structure. An example of such diseases is Parkinson’s Disease (PD) where α-synuclein (αS) aggregates appear to be the cell-toxic agent.^1–3^ It has been proposed that such amyloids may form as an immune response to an infection, rendering pathogens ineffective by entrapping them^4^. While evidence for this microbial protection hypothesis^5–7^ is limited, correlations have been observed, for instance, between falling ill with COVID-19 and outbreaks of PD^8^. Because of these correlations, and SARS-COV-2 promoted αS amyloid formation *in vitro*^9–11^, we have proposed earlier^12^ that amyloidogenic regions on SARS-COV-2 proteins can enhance aggregation of αS. In these previous studies we investigated the interactions of the fragment SFYVYSRVK (SK9) of the Envelope protein with αS. We found that SK9 alters the ensemble of αS monomer conformations toward such that are more aggregation prone, preferentially raising the likelihood for forming the rod polymorph^13,14^. By themselves can these preliminary investigations be criticized for relying on an isolated short SARS-COV-2 protein fragment, instead of the complete viral protein where the spatial environment of the segment may change its ability to cross-seed αS. A related question is why the suspected health effects are not seen after infections by other Corona viruses that have homologous segments. However, it has been recently shown that such short fragments, derived by enzymatic cleavage of SARS-COV-2 proteins, exist in humans, and that in vitro they form amyloids^15^. Of special interest is the segment 194–203, (FKNIDGYFKI), of the spike protein as it is unique for the SARS-COV-2 virus, and as it is cleaved from the Spike protein by the enzyme neutrophil elastase. This enzyme is released from neutrophils during acute inflammation, as, for instance, caused by the infection. Motivated by these experimental investigations we have researched with extensive molecular dynamics simulations the effect of presence of the SARS-COV-2 spike protein segment FKNIDGYFKI, which we call in the following FI10, on αS monomers and fibrils. Our goal is to demonstrate that our previous results are not an artifact of the SK9 fragment, but also seen for the likely more disease-relevant and SARS-COV-2 specific FI10. For this purpose, we compare the effect that the two fragments have on αS monomers and fibrils to establish commonalities and differences. Our results let us to conjecture that viral infections raise the risk for amyloidosis when, as often in the case of SARS-COV-2, the infection causes pronounced inflammation and release of enzymes that can cleave viral proteins into in some cases amyloidogenic fragments which can cross-seed human proteins.

## II. MATERIALS AND METHODS

### A. System Preparation

Using molecular dynamics (MD) simulations we investigate how SARS-COV-2 proteins alter αS aggregation. For this purpose, we compare the effect of the ten-residue segment FKNIDGYFKI (FI10) of the Spike protein on the conformational ensemble of αS monomers and the stability of αS fibrils with that induced by the nine-residue segment SFYVYSRVK (SK9) of the Envelope protein studied by us in earlier work.^12,16^ As start conformation for the αS we used either the model of micelle-bound αS deposited in the Protein Data Bank (PDB) under PDB ID: 1XQ8, or a randomized and fully-extended conformation derived by heating up the PDB conformation simulating it at 500 K for 5 ns^12^. For the fibril models we extracted decamers from the cryo-EM structures deposited in the PDB under identifiers 6CU7 (rod-like architecture) and 6CU8 (twister-like architecture).^13^ In order to be consistent with earlier work^12^, we add again at the N- and C-terminal a NH^3+^ and a COO^−^ group for both monomer and fibril models.

Simulations starting from the above described αS monomer and fibril models described above serve as control for simulations where in addition also either SK9 or FI10 peptides are present. Following Nyström and Hammarström^15^ the FI10 viral fragment was derived from the structure of the SARS-COV-2 Spike protein (PDB ID: 6VXX) by extracting the segment of residues F194-I203, and capping the N- and C-terminal of this peptide by a NH^3+^ and -COO^-^ group. Docking FI10 segments in a ratio of 1:1 with either monomer or fibrils at binding sites predicted by AutoDock Vina^17^ or HADDOCK^18,19^ we obtain the initial conformations for our simulations, shown in **Figure 1**. Note that the viral protein segments are not tethered to these initial positions but move freely throughout the simulations and may detach from the αS chains. In a few cases we extended or repeated previous simulations of SK9 interacting with αS monomers or fibrils. In these instances we followed the preparation described in our earlier work.^12,16^

**FIGURE 1:**
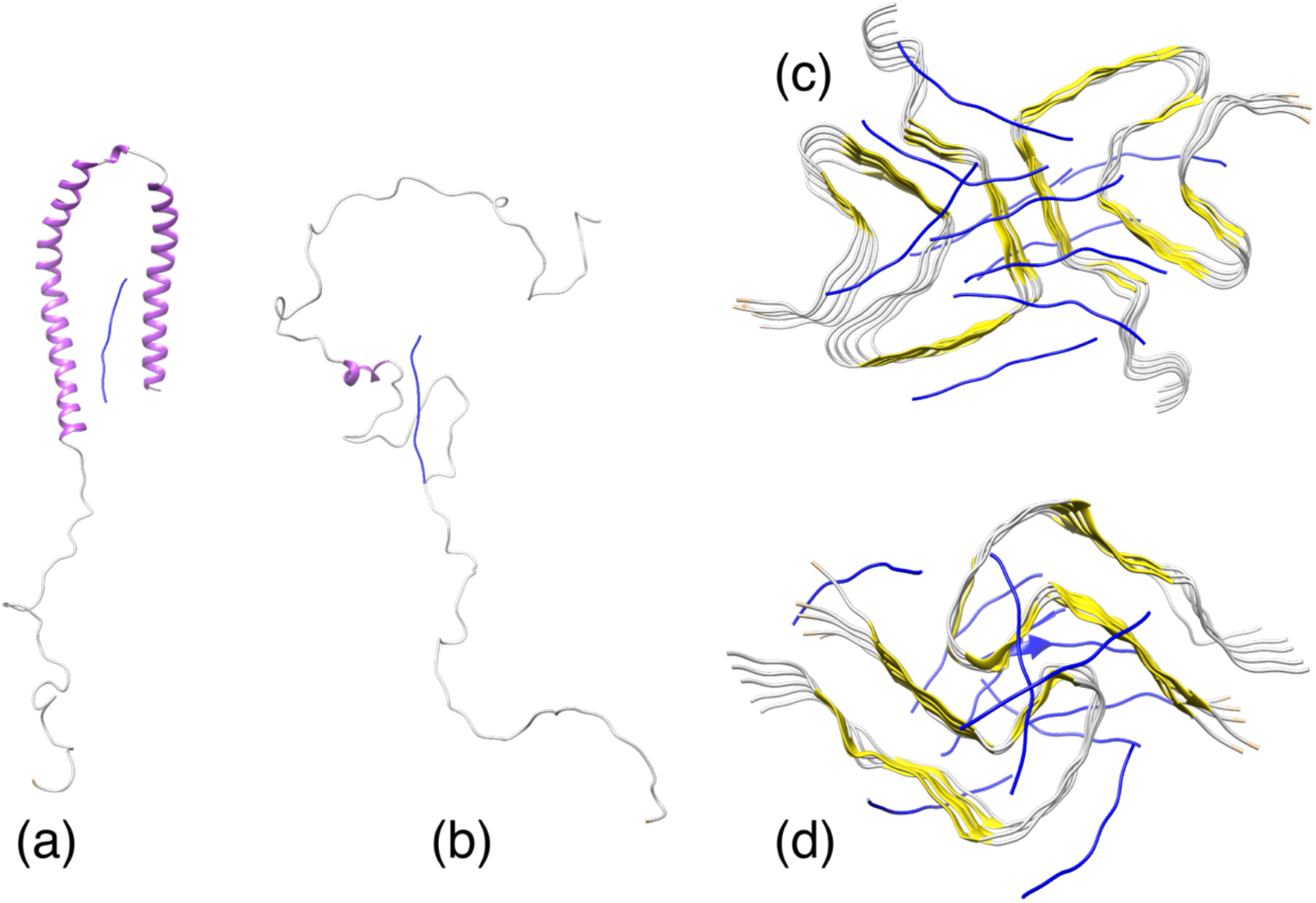
Start configurations of αS with initial binding positions of FI10: (a) monomer (PDB ID: 1XQ8), (b) heat-extended monomer, (c) rod-polymorph (PDB ID: 6CU7) fibril, and (d) twister-polymorph (PDB ID: 6CU8) fibril.

### B. General Simulation Protocol

Following our earlier work we simulate the monomer systems with the a99SB-disp all-atom force-field^20^ in combination the corresponding a99SB-disp water model, whereas the fibrils simulations rely on the CHARMM 36m all-atom force-field^21^with TIP3P explicit water^22^. Both force-field are implemented in the GROMACS package 2022 package^23^ used for our simulations. Hydrogen atoms were added with the *pdb2gmx* module of the GROMACS suite^23^. The start configurations for each system were placed in the center of a cubic box with periodic boundary conditions, with a minimum distance of 15 Å between the solute and the edge of the box. The boxes were filled with water molecules, and counterions added to neutralize the system. Na^+^ and Cl^-^ ions are at a physiological ion concentration of 150 mM NaCl. The number of water molecules and the total number of atoms are listed in **Table I**. The energy of each system was minimized by steepest decent for up to 50,000 steps, and afterwards the system was equilibrated at 310K for 200 ps at constant volume and an additional 200ps at constant pressure (1 atm), constraining the positions of heavy atoms with a force constant of 1000 kJ mol^-1^ nm^-2^.

**Table I:** Simulated systems

| System Description | Atoms | Water Molecules | Independent Trajectories | Simulation Length (ns) | Total Sampling (ns) |
| --- | --- | --- | --- | --- | --- |
| **Monomer (PDB ID: 1XQ8)** |  |  |  |  |  |
| Control | 337,169 | 83,670 | 3 | 1,000 | 3,000 |
| w/ SK9 | 336,308 | 83,414 | 3 | 1,000 | 3,000 |
| w/ FI10 | 336,359 | 83,423 | 3 | 1,000 | 3,000 |
| **Monomer (EXTENDED)** |  |  |  |  |  |
| Control | 336,177 | 83,422 | 2 | 1,000 | 2,000 |
| w/ SK9 | 336,132 | 83,370 | 2 | 1,000 | 2,000 |
| w/ FI10 | 336,283 | 83,404 | 2 | 1,000 | 2,000 |
| **Rod Fibril Model (PDB ID: 6CU7)** |  |  |  |  |  |
| Control | 183,222 | 58,098 | 3 | 200 | 600 |
| w/ SK9 | 183,383 | 57,585 | 3 | 200 | 600 |
| w/ FI10 | 215,356 | 68,190 | 3 | 200 | 600 |
| **Twister Fibril Model (PDB ID: 6CU8)** |  |  |  |  |  |
| Control | 214,717 | 69,447 | 3 | 200 | 600 |
| w/ SK9 | 177,583 | 56,525 | 3 | 200 | 600 |
| w/ FI10 | 215,101 | 68,975 | 3 | 200 | 600 |

Simulations started from the above generated initial conformations (shown in **Figure 1**). A constant temperature (310 K) was maintained by a v-rescale thermostat^24^ with a coupling constant of 0.1 ps, at and pressure was kept constant 1 atm by using the Parrinello-Rahman barostat^25^ with a pressure relaxation time of 2 ps. By using the SETTLE algorithm^26^ to keep water molecules rigid, and restraining protein bonds involving hydrogen atoms to their equilibrium length with the LINCS algorithm^27^, we were able to use a time step of 2 fs for integrating the equations of motion. Since we employed periodic boundary conditions, we computed the long-range electrostatic interactions with the particle-mesh Ewald (PME) technique, using a real-space cutoff of 12 Å and a Fourier grid spacing of 1.6 Å. Short-range van der Waal interactions were truncated at 12 Å, with smoothing starting at 10.5 Å. For each system, we follow three independent trajectories differing by their initial velocity distributions. The length of the various trajectories is also listed in **Table I**. PDB-files of start and final configurations are available as **supplemental material**.

### C. Trajectory Analysis

The molecular dynamics trajectories are analyzed with the GROMACS tools^23^, VMD^28^ and MDTraj software^29^. For visualization of conformations we use VMD^28^. GROMACS tools are used to calculate root-mean-square deviation (RMSD), root-mean-square-fluctuation (RMSF), radius of gyration (RGY) and solvent accessible surface area (SASA), the later quantity relying on a spherical probe of 1.4 Å radius. Contact frequencies are calculated using VMD and the MDTraj software, defining contacts by a cutoff 4.5 Å in the closest distance between heavy atoms in a residue pair. The residue-wise secondary structure propensity is calculated in VMD by the STRIDE algorithm^30^. Helicity and sheetness are defined as the percentage of residues with alpha-helical (i.e., assigned a “H” by STRIDE) or beta-strand (an “E” for extended configuration or a “B” for an isolated bridge) secondary structure in a frame of the trajectory analyzed.

## II. RESULTS AND DISCUSSION

### A. Monomer simulations

In previous work,^12^ we established that interaction with the SARS-COV-2 Envelope protein fragment SK9 moves the ensemble of αS chain structures toward more extended and aggregation prone ones. We also argued there that these changes favor assembly into rod-like amyloids. In the present work we compare these results with that of simulations where we consider instead interaction of αS chains with the fragment FI10 of the Spike protein as this fragment is unique for the SARS-COV-2 virus and is cleaved from the Spike protein under disease conditions. We remark that our results are averages over six runs, with three starting from a micelle-bound and three from an extended αS conformations (see method section). This is because we saw few differences between the two sets of runs in the last 50 ns of trajectories considered by us.

We first compare the strength with which both viral protein fragments bind to αS chains. In the start conformations SK9 has 27 contacts with the αS monomer; and FI10 forms 22 contacts. The diffusive motion of the fragments in relation to the αS chain leads to a loss of many of these contacts (on average there are for SK9 nine of the original contacts left in the final 50 ns of the trajectory, and none for FI10), but some are replaced by newly formed contacts. Consequently, the total number of binding contacts decreases more slowly, and the average number of binding contacts over the last 50 ns are 21(10) for SK9, and 5(3) for FI10. On the other hand, when calculated by the method of Ref.^31^, we find binding energies of -19(4) kJ/mol for SK9 and -20(5) kJ/mol for FI10, that despite the lower number of contacts suggests a tighter binding of FI10 to the αS monomer. **Figure 2a** shows that for FI10 the persisting contacts are located in the segment A90–I112 in the C-terminus, while SK9 binds mainly to residues K21-E35 at the N-terminus, the NAC region of residues E61 to A78 and the disorganized C-terminus starting residue D119. We find that, because of these contacts, FI10 increases the flexibility of residues over the whole αS chain, see **Figure 2b**, but especially leads to an increased flexibility for the segments of residues E46 -V70 and T75-V95, i.e., the NAC and preNAC region. On the other hand, for SK9 the increase in flexibility is more narrowly restricted to the short segments where it binds strongly to the αS monomer. We remark that in the much longer simulations of Ref.^12^ presence of SK9 finally also results in a higher flexibility of residues over almost the whole αS chain.

**Figure 2:**
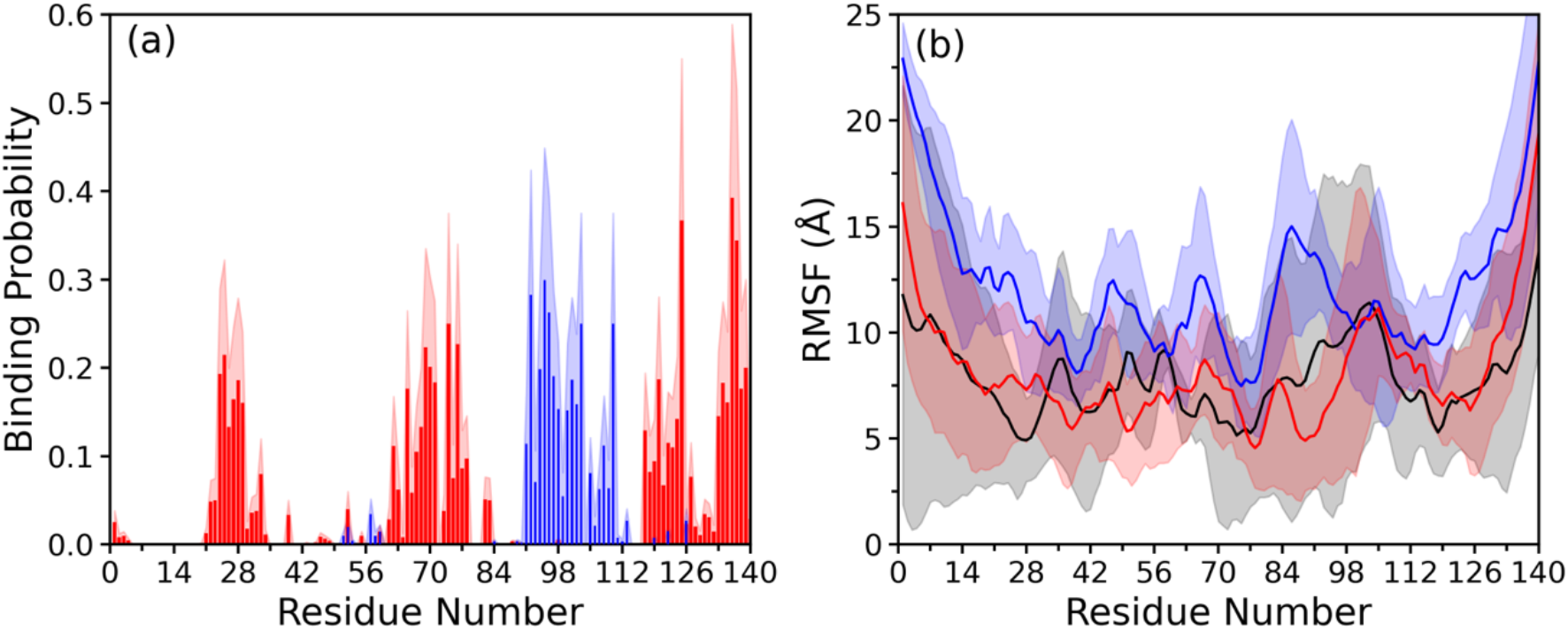
(a) The residue-wise normalized binding probability of the FI10 (blue) or the SK9 (red) segment to αS monomer. (b) Residue-wise mean square fluctuation (RMSF; measured in Å) obtained from αS monomer simulations in the absence (in black) and in the presence of the SK9 (red) or the FI10 peptide (blue). The RMSF values are calculated with reference to the experimentally resolved αS monomer (PDB ID: 1XQ8) considering only backbone atoms. Data are averaged over the final 50 ns of each trajectory. Shaded region represents the standard deviation of the average values defining the corresponding curves.

The net-effect of this higher flexibility is that the αS monomers are more extended and strand-like when in presence of SK9 or FI10. This can be seen in **Figure 3a** where we show the distribution of the radius of gyration (R_g_), a measured for the compactness of conformations, as obtained in the last 50 ns, and averaged over all three trajectories. Interaction with the viral protein fragments shifts the ensemble towards less compact conformations. This effect is more pronounced for FI10 than for SK9 as can be also seen in the corresponding averages listed in **Table II**. The distribution of solvent-accessible surface area (SASA) in **Figure 3b** support the much larger effect that FI10 has on the ensemble of αS monomer conformations. Correspondingly we find a much lower number of contacts between residues within the αS monomers when in presence of FI10 than in the control of in presence of SK9, see **Table II**. Hence, presence of FI10 causes a pronounced shift in the ensemble of monomer conformations toward more extended, solvent-exposed and looser-packed conformations that in this magnitude is for SK9 only observed over much larger time scales (3 μs instead of 1μs), see Ref.^12^.

**Table II:**
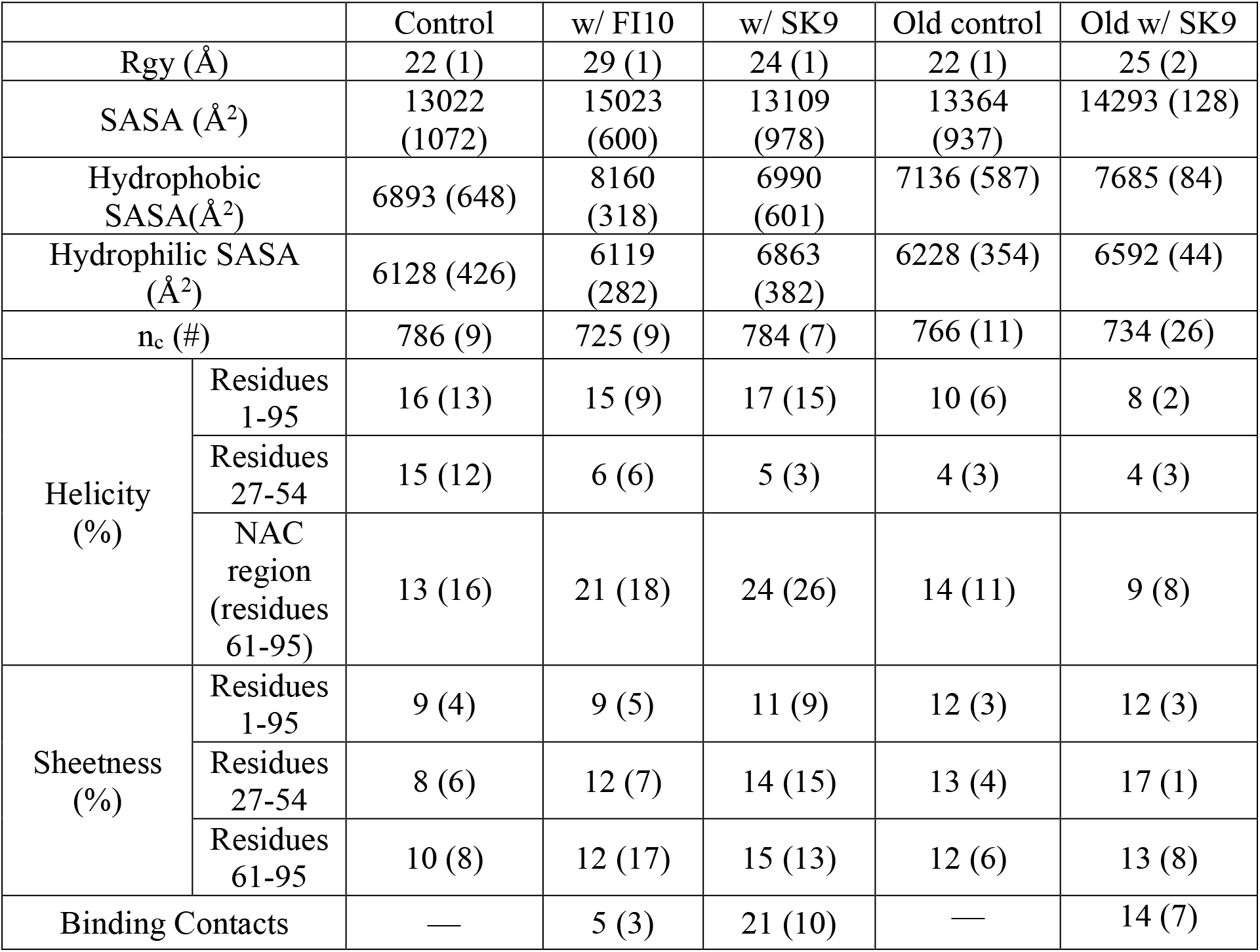
for control, SK9, FI10 average RGY, SASA, nc, helicity, sheetness (whole range, 27-54,61-95), and binding contacts. Averages over last 50ns. Data from a previous study published in Ref12 added separately for reference (data calculated over last 1.5 μs of 3μs trajectories).

|  |  | Control | w/ FI10 | w/ SK9 | Old control | Old w/ SK9 |
| --- | --- | --- | --- | --- | --- | --- |
| Rgy (Å) |  | 22 (1) | 29 (1) | 24 (1) | 22 (1) | 25 (2) |
| SASA (Å<sup>2</sup>) |  | 13022<br>(1072) | 15023<br>(600) | 13109<br>(978) | 13364<br>(937) | 14293 (128) |
| Hydrophobic SASA(Å<sup>2</sup>) |  | 6893 (648) | 8160<br>(318) | 6990<br>(601) | 7136 (587) | 7685 (84) |
| Hydrophilic SASA (Å<sup>2</sup>) |  | 6128 (426) | 6119<br>(282) | 6863<br>(382) | 6228 (354) | 6592 (44) |
| n<sub>c</sub> (#) |  | 786 (9) | 725 (9) | 784 (7) | 766 (11) | 734 (26) |
| Helicity (%) | Residues 1-95 | 16 (13) | 15 (9) | 17 (15) | 10 (6) | 8 (2) |
|  | Residues 27-54 | 15 (12) | 6 (6) | 5 (3) | 4 (3) | 4 (3) |
|  | NAC region (residues 61-95) | 13 (16) | 21 (18) | 24 (26) | 14 (11) | 9 (8) |
| Sheetness (%) | Residues 1-95 | 9 (4) | 9 (5) | 11 (9) | 12 (3) | 12 (3) |
|  | Residues 27-54 | 8 (6) | 12 (7) | 14 (15) | 13 (4) | 17 (1) |
|  | Residues 61-95 | 10 (8) | 12 (17) | 15 (13) | 12 (6) | 13 (8) |
| Binding Contacts |  | — | 5 (3) | 21 (10) | — | 14 (7) |

**Figure 3:**
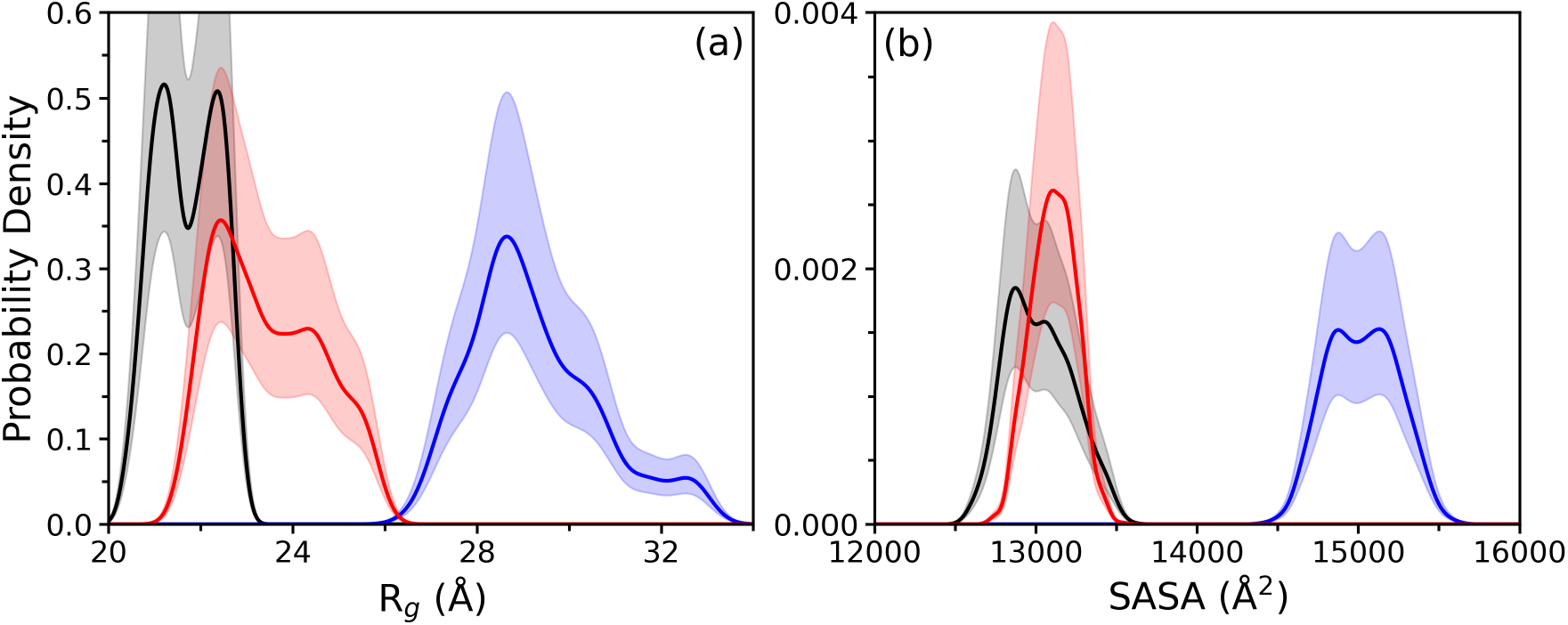
Probability density of (a) radius of gyration (R_g_) and (b) solvent accessible surface area (SASA) of αS monomers in absence (black) and presence of the SK9 (red) and FI10 (blue) peptide. Data are averaged over the final 50 ns of each trajectory. Shaded region represents the standard deviation of the average values defining the corresponding curves.

The decrease in the average frequency of helicity and the corresponding gain in the frequency of sheetness seen in **Table II** indicates that the resulting conformations are more aggregation prone. The change in secondary structure is most pronounced for the N-terminal segment of residues A27–T54. Most of the known familial mutations are located in this segment assumed to be important in the early stages of αS aggregation.^32–36^ In both cases is the helicity reduced from about 15% to 5%-6%, while at the same time the sheetness increases from about 8% to 12%-14% and is slightly higher for FI10. Interestingly, we find in the NAC region of residues E61-V95 an increase in both helicity and sheetness.

As already observed in Ref.^12^, the shift in the αS monomer ensemble seems to be not just one toward more aggregation-prone conformations, but also favoring assembly into rod-like fibrils over such with a twister-like structure. In the rod fibril polymorph (see **Figure 1**) the segment E46-A56 forms the interface between the two protofibrils while for the twister polymorph the interface is formed by the region G68-A78. In our simulations of the αS monomer we find for the segment E46–A56 the solvent accessible surface area (SASA) raised by 90 Å^2^ in the presence of SK9 (from 1334 (111) Å^2^ to 1422 (108) Å^2^), and by about 104 Å^2^ in the presence of FI10 (to 1437 (100) Å^2^). The difference in SASA arises in the case of SK9 almost equally from hydrophilic (48 Å^2^) and hydrophobic residues (42 Å^2^), while for FI10 the difference is mostly due to more exposed hydrophobic residues, (65 Å^2^ vs. 39 Å^2^ for hydrophilic residues). On the other hand, for the segment G68-A78 the SASA values raise only marginally from 1286 (122) Å^2^ for the control to 1303 (85) in presence of SK9, and even decrease in presence of FI10 to 1268 (140) Å^2^. This decrease is from hydrophilic residues (-34 Å^2^) while hydrophobic residues are slightly more exposed by about 16 Å^2^. For SK9 arises the difference in SASA almost equally from hydrophilic and hydrophobic residues. Hence, presence of both SK9 and FI10 leads to an exposure of solvent accessible surface (especially of hydrophobic residues) for the segment E46-A56 that is much lower or not existing for the segment G68-A78. As a consequence, association of αS chains is favored in both cases more at the segment forming in the rod-like fibril the inter-protofilament interface than at the corresponding segment in the twister-like fibril. This effect is again especially pronounced in the case of FI10 than for SK9, with FI10 even decreasing the SASA values of the segment G68-A78. On the other hand, even in the longer simulations of SK9 interacting with As of Ref.^12^ the SASA values of 1444(28) Å^2^ for the segment E46–A56 (as measured over the last 1.5 μs of the 3 μs) barely match the values measured in the FI10 simulation already much earlier at 1μs, while for G68-A78 they further increase to 1335(46) Å^2^.

Hence, our monomer data confirm that alteration of the ensemble of αS monomer conformations is not restricted to the SK9 segment studied earlier by us. The net effect of the interaction between SK9 or FI10 and αS monomers is in both cases a shift toward monomer conformations that have a higher propensity to aggregate. The signal for this shift is stronger for the Spike protein fragment FI10 which is unique for SARS-COV-2. Even more than SK9 does FI10 leads to more extended conformations with more exposed solvent accessible surface, with an exposure of hydrophobic residues favoring association into rod-like over twister-like fibrils.

### B. Fibril simulations

Another way by which SARS-COV-2 proteins can promote amyloid formation is by raising the stability of fibril polymorphs. For this reason, we have probed in a second set of molecular dynamics simulations how SK9 or FI10 change the stability of the two most common αS fibril structures, the rod and the twister polymorphs (see **Figure 1)**. As with the monomer simulations, we compare for the two fibril polymorphs such trajectories, where SK9 or FI10 fragments bind initially to the αS chains, with the corresponding ones where the viral protein fragments are absent.

While for the rod model only 60 residues (L38-K97) out of the 140 amino acids in αS have been resolved, and only 41 residues (K43–E83) for the twister model, we decided against adding the unresolved regions as predicted by homology modeling. This is because we found in previous work^12^ no qualitative differences in the behavior of the original PDB models and the ones with extended αS chains.

Initially, the SK9 peptides form about 27 contacts per chain the 6CU7 rod fibril, and the FI10 peptides form on average 33 contacts per chain. These numbers decrease quickly in the first nanosecond, and more slowly afterwards, but stay constant over the last 50 ns at 13(1) contacts per chain for SK9, and 16 (1) contacts for FI10, i.e., the number of contacts decreased by about 50% in both cases. Note that not all of these contacts appear in the start configuration: neither the SK9 nor the FI10 fragments are tethered to the fibril, and binding contacts can both form and dissolve. The interaction between FI10 peptides and the 6CU7 rod fibril is stronger than it is between SK9 fragments and fibril, with the residue-wise distribution of contacts much broader for FI10 than for SK9 where the binding sites are localized to segments of residues H50–K60 and V71–Q79, see **Figure 4a**.

**Figure 4:**
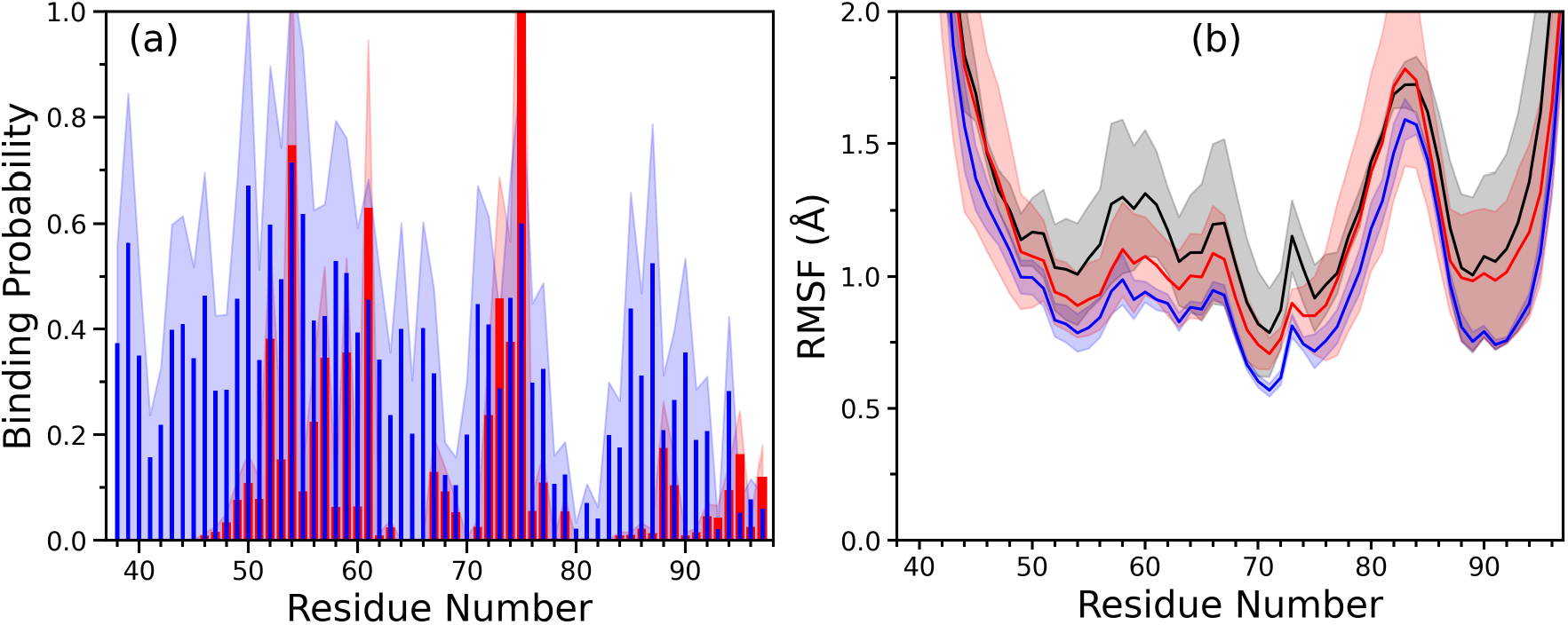
The residue-wise normalized binding probability of the SK9 (red) and the FI10 (blue) peptides to αS chains in the rod-like fibril model 6CU7 is shown in (a), while we display in (b) the corresponding residue-wise root-means-square fluctuations (RMSF) for αS chains (SK9: red, FI10: blue, control: black). Data are averaged over the final 50 ns of each trajectory and shaded regions represent the standard deviation of the average values defining the curves.

The binding with the viral protein fragment leads in both cases to a reduced flexibility of the αS chains, see the lower values of the residue-wise root mean square fluctuation (RMSF) in **Figure 4b**. While the overall form of all three RMSF curves is similar, residues in the 6CU7 rod fibril bound to SK9 or FI10 have lower RMSF values than the control, i.e., are less flexible, and the decrease in flexibility is closely correlated with the binding probabilities for the viral protein fragments to the αS chains in **Figure 4a**. While the RMSF distribution indicates that both SK9 and FI10 stabilize the rod-like fibril, the effect more pronounced for FI10. Note especially the strong additional stabilization of chain geometry at the inter-protofilament interface (E46–A56), but also the stabilization of the C-terminal residues G86–K96 that is not seen for SK9.

Corresponding to the reduced chain flexibility are the final conformations less perturbed in presence of both viral protein fragments than for the control. Only minor changes in the stacking and packing pattern are observed, and these help to preserve the overall fibril topology, see the representative final conformations in **Figure 5a-c**. As the structural changes are mainly in the individual chains and not the overall topology of the fibril, we show in **Figure 5d** the chain root-mean-square deviation (RMSD) as function of time. Here, the RMSD calculated separately for each chain, taking only backbone atoms into account, and averaged over all ten chains in the fibril model. While the differences to the control are small, the RMSD is slightly lower when SK9 or FI10 bind to the 6CU7 rod fibril. The stabilization by SK9 and FI10 is in both cases connected with an increase in intrachain contacts (from 2410 (30) in the control to 2590 (50) for SK9 and 2533 (6) for FI10). For FI10, the increase in intrachain contacts is correlated with one in stacking contacts between layers of chains and leads to a reduced solvent accessible surface area (29400(600) Å^2^ vs 29900(500) Å^2^ in the control), with the difference mostly from a reduction of hydrophobic SASA (15800(300) Å^2^ vs 16200(300) Å^2^ in the control). However, in presence of SK9 the SASA is increased to 30500(700) Å^2^, likely because of a lower number of packing contacts that ease separation of the two protofibrils and leads to increased exposure of hydrophilic residues, see **Table III**.

**Table III:** Mean values of hydrophobic SASA, hydrophilic SASA, total SASA, number of intrachain, stacking and packing contacts, and frequency of native interchain and intrachain contacts. All values are averages over the final 50 ns of all trajectories of the respective systems.

| system | hydrophobic SASA (Å<sup>2</sup>) | hydrophilic SASA (Å<sup>2</sup>) | total SASA (Å<sup>2</sup>) | intrachain contacts | stacking contacts | packing contacts | % of native interchain contacts | % of native intrachain contacts |
| --- | --- | --- | --- | --- | --- | --- | --- | --- |
| Rod 6CU7 control | 16200 (300) | 13700 (100) | 29900 (500) | 2410 (30) | 1130 (20) | 97 (6) | 51 (4) | 40 (1) |
| Rod 6CU7 w/ SK9 | 16300 (500) | 14200 (300) | 30500 (700) | 2590 (50) | 1130 (30) | 89 (5) | 54 (5) | 40 (2) |
| Rod<br>6CU7<br>w/<br>FI10 | 15800 (300) | 13600<br>(300) | 29400<br>(600) | 2533 (6) | 1150<br>(20) | 100<br>(20) | 38 (6) | 40 (1) |
| Twister<br>6CU8<br>control | 10700 (400) | 10700<br>(400) | 21400<br>(800) | 1670 (10) | 700<br>(20) | 125 (3) | 43 (3) | 42 (1) |
| Twister<br>6CU8<br>w/ SK9 | 10600 (200) | 11000<br>(150) | 21600<br>(400) | 1642 (1) | 720<br>(10) | 112 (5) | 41 (1) | 40 (0) |
| Twister<br>6CU8<br>w/<br>FI10 | 10800 (100) | 10900<br>(300) | 21700<br>(200) | 1644 (7) | 719 (4) | 119 (7) | 49 (1) | 41 (1) |

**Figure 5:**
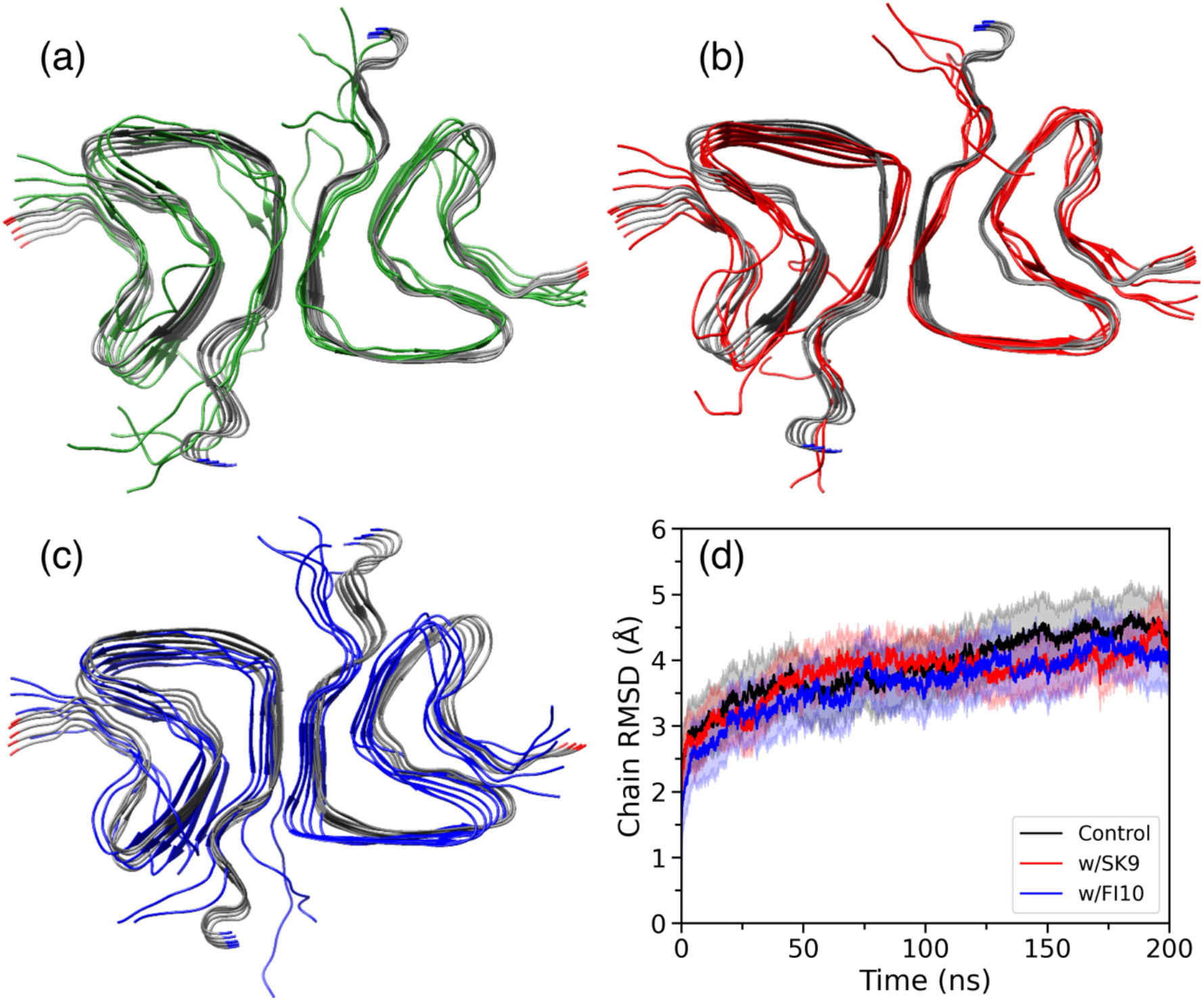
Representative conformations after 200 ns simulations of the rod-like αS fibril model 6CU7 in (a) absence and presence of (b) SK9 or (c) FI10. For comparison we show these conformations overlayed over the start conformation. The chain rmsd as function of time is shown for the three cases in (d), with shaded region representing the standard deviations.

On the other hand, in the 6CU8 twister fibril, per αS chain the SK9 peptides form about 31 contacts and the FI10 peptides 27 contacts. The number of contacts decreases in both cases but stays constant over the last 50 ns at 24 (1) contacts per chain for SK9, and 16 (1) contacts for FI10, i.e., the number of contacts decreased by about 20% for SK9 and 40% for FI10. As in the case of the rod-fibril simulations, neither the SK9 nor the FI10 fragment is tethered to the fibril and binding contacts may form and dissolve. Unlike for the rod-fibril, the residue-wise distribution of contacts between viral protein fragment and the αS chains in **Figure 6a** is broad for both SK9 and FI10 and covers the whole chain, with a peak for SK9 around residues H50-K58. This region of increased binding probability coincidences with the sole part of the αS chain, the segment of residues H50 to D65, where presence of SK9 leads to a pronounced decrease in flexibility that is not seen for FI10. Otherwise, few differences are seen in the residue-wise RMSD distribution when comparing the ones for SK9 and FI10, see **Figure 6b**. Note especially that the flexibility of the region G68-A78, forming the interface between the two protofibrils in the twister polymorph, is equally reduced in presence of either of the two peptides. However, the decrease in flexibility caused by either of the two peptides appears in general to be smaller than in the case of the rod-like fibril.

**Figure 6:**
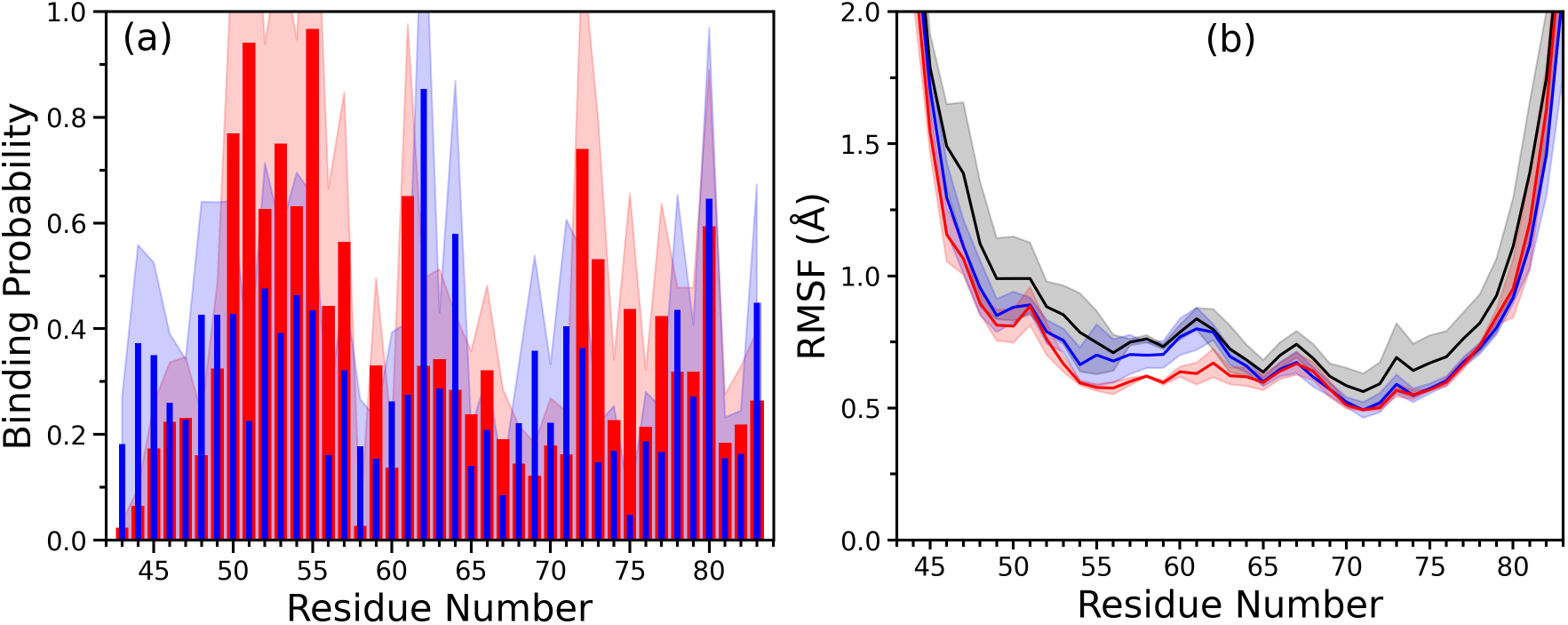
The residue-wise normalized binding probability of the SK9 segment (red) and the FI10 segment (blue) to αS chains in the twister-like fibril 6CU8 is shown in (a), while in (b) we show for this model the residue-wise RMSF for the αS chains in presence of SK9 (red), FI10 (blue) and in the control (black). Data are averaged over the final 50 ns of each trajectory and shaded region represents the standard deviation.

While as for the rod-like fibril the final conformations are in presence of SK9 or FI10 less perturbed than the control, see the representative final conformations in **Figure 7a-c**, we see for the twister polymorph also a clear difference in the chain RMSD curves between SK9 and FI10, see **Figure 7d**. While the values for FI10 are comparable to the control, they are for SK9 substantially lower, indicating increased stability in presence of SK9 peptides, but not in presence of FI10 fragments. This reduced RMSD likely result from the binding of SK9 to the segment H50-K58 and the induced reduction in flexibility for the segment H50-D65. However, the various quantities shown in **Table III** do not support this picture and show only marginal differences between SK9 and FI10. While the percentage of remaining native intrachain contacts (i.e., such already present in the start conformation) changes little between rod and twister fibril, the number of interchain contacts decreases from about 50% to 40% for the SK9 complex and in the control, but it is 10% higher for FI10. This is not surprising as FI10 has the broad residue-wise binding distribution, influencing more strongly interchain binding. For the twister this leads in the case of FI10 also to a slightly increased number of stacking contacts and similar decreased number of packing contacts. However, consistent with the visual inspection of the final conformations in **Figures 7a-c** do the various quantities in **Table III** indicate that presence of SK9 or FI10 stabilizes the twister fibril only marginally, with little differences between the two viral protein fragments. Hence, presence of the viral protein fragments affects the twister-like fibril very different from the rod-like fibril where we saw a clearer signal indicating stabilization by FI10, and to a lesser degree also by SK9

**Figure 7:**
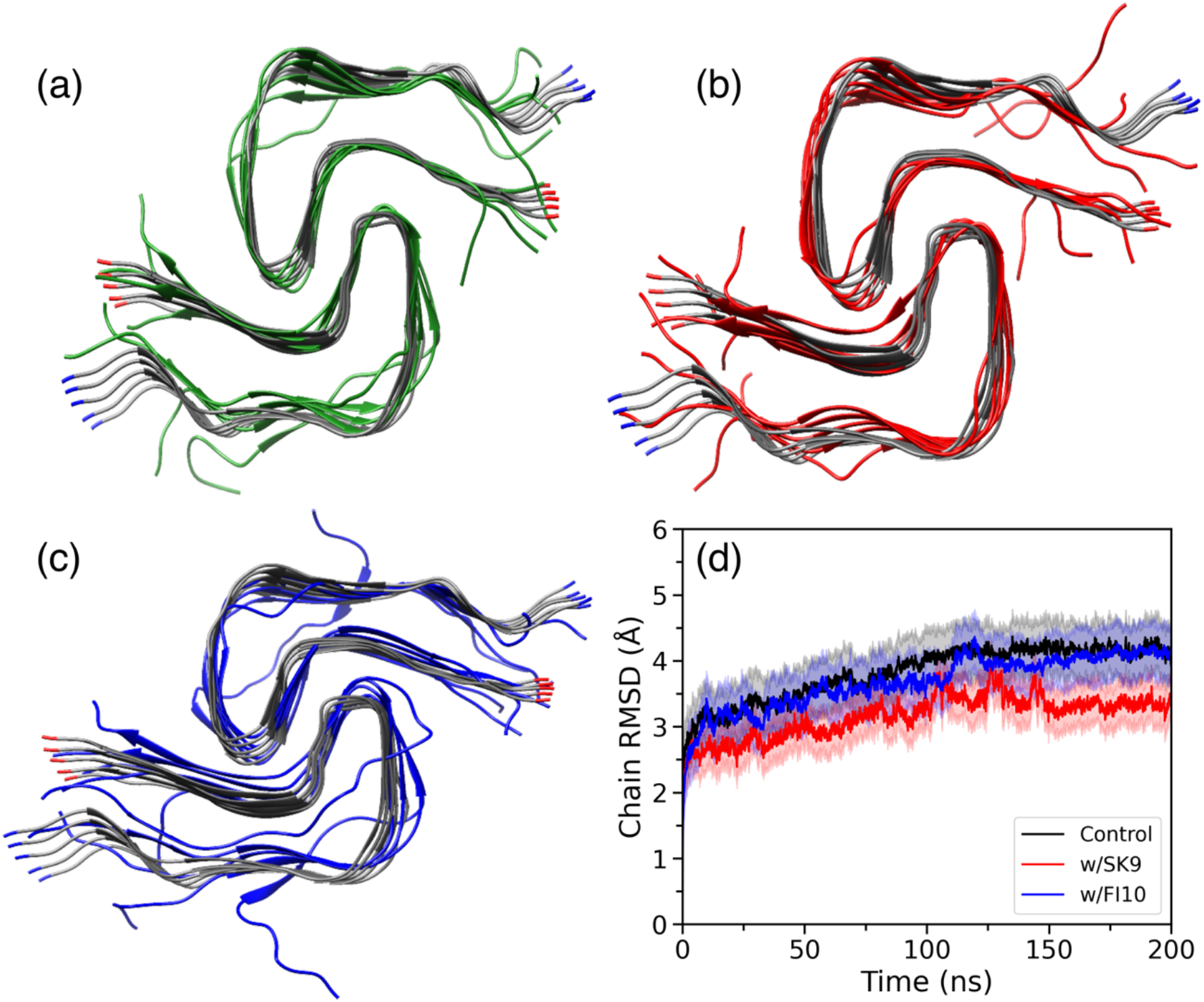
Representative conformations after 200 ns simulations of the rod-like αS fibril model 6CU8 in (a) absence and presence of (b) SK9 or (c) FI10. For comparison we show these conformations overlayed over the start conformation. The chain rmsd as function of time is shown for the three cases in (d), with shaded region representing the standard deviation.

## III. CONCLUSIONS

Using molecular dynamics simulations, we have studied the effect of two protein fragments of the SARS-COV-2 virus on the structural ensemble of αS monomer, and on the stability of αS fibrils. One virus protein fragment is from the Envelope protein (SK9) and one from the Spike protein (FI10). Both bind to the αS monomer, and induce the shift toward more aggregation-prone conformations that we also saw in our previous studies, not only of αS but also with Serum Amyloid A and amylin.^16,37^ This effect is stronger for FI10, a fragment that is unique for SARS-COV-2 and cleaved from the Spike protein by enzymes released during pronounced inflammation.^15^ Especially in the case of FI10 leads the shift in the ensemble of αS chains to conformations with preference for assembly into rod-like fibrils. We also find again the earlier observed stabilization of fibrils. This effect is differential. While both SK9 and FI10 alter only marginally the stability of the twister polymorph, the stability of rod-like fibrils is clearly enhanced, with the effect more pronounced for FI10 than for SK9.

Hence, while the time scales of our simulations are small compared to that of amyloid formations, our data indicate already that SARS-COV-2 protein fragments can induce amyloid formation of αS (implicated in Parkinson Disease), and that this effect is differentially, i.e., favoring certain polymorphs, in our case rod-like αS fibrils over fibrils with a twister-like chain structure. However, the relation between αS polymorphs and disease phenotype has not been established.

While fibrils seeded with lysates from brain tissue of Parkinson Disease patients have rod-like structures,^38^ this may be an artefact of the growth conditions. On the other hand, the clustering of familial mutations associated with Parkinson disease in the preNAC region, de-stabilizing the rod-architecture but not twister-like conformations, rather points to twister polymorphs as disease-causing agent.^13^ Hence, it is not clear whether our observation points to a mechanism that may explain the correlation between SARS-COV-2 infections and early onset of Parkinson Disease. Nevertheless, our results add to an emerging picture of SARS-COV-2 protein fragments (cleaved during infection-cause inflammation) cross-seeding amyloidogenic human proteins, potentially triggering the onset of the associate amyloid diseases.

## Supporting information

Start and final conformations for all trajectories

## SUPPLEMENTARY MATERIAL

The Supplementary Material is available free of charge at URL

- Start and final configurations of all trajectories in PDB format as text files in a compressed folder

## ACKNOWLEDGMENT

The simulations in this work were done using the SCHOONER cluster of the University of Oklahoma, XSEDE resources allocated under grant MCB160005 (National Science Foundation), and TACC resources allocated under grant under grant MCB20016 (National Science Foundation). We acknowledge financial support from the National Institutes of Health under grant GM120634. We thank Shailesh K. Panday for help at an early stage of this study with setting up and running simulations.

## AUTHOR DECLARATIONS

### Author Contributions

**Andrew D. Chesney:** Formal analysis (equal); Investigation (equal); Visualization (lead); Writing – original draft (equal); Writing – review and editing (equal); **Buddhadev Maiti:** Formal analysis (equal); Investigation (equal); Writing – original draft (equal); Writing – review and editing (equal); **Ulrich H.E. Hansmann:** Conceptualization (lead); Funding acquisition (lead); Resources (lead); Supervision (lead); Writing – original draft (equal); Writing – review and editing (equal).

### DATA AVAILABILITY

The data that support the findings of this study are available from the corresponding author upon reasonable request.

